# Angiogenesis Guided by *Bombyx mori* Silk Proteins is Molecular Weight-Dependent

**DOI:** 10.64898/2026.06.28.735127

**Authors:** Tiandong Li, Jian He, Jiajin Qian, Yeyuan Wang, Jingchen Sun, Doudou Hu

## Abstract

Silk proteins, including sericin and fibroin, are natural biopolymers with broad applications in tissue engineering where angiogenesis plays an essential role. However, the pro-angiogenic effects of silk proteins with varying molecular weights (MWs) remain poorly understood. Here, silk proteins with MW distributions at 40-180 kDa or less than 25 kDa were obtained through alkaline hydrolysis to evaluate their effects on angiogenesis. Structurally, reducing MW induced a conformational transition in silk proteins, accompanied by a striking morphological shift in sericin from nanofibers to nanoparticles. Functionally, high-MW sericin (SSH) suppressed, whereas low-MW sericin (SSL) and both high- and low-MW silk fibroin (SFH/SFL) directly promoted endothelial angiogenic activity. Transcriptomic analysis revealed that angiogenesis-related genes such as *Id1* and *Smad6/9* may underlie the angiostatic effects of SSH. Notably, both SSH and SSL enhanced angiogenesis indirectly via macrophages; however, SSH induced mixed M1/M2-like polarization, while SSL preferentially drove an M2-like phenotype. In a subcutaneous implantation model, SSH promoted angiogenesis but yielded vessels with weak integrity and increased fibrosis, whereas SSL enhanced angiogenesis with improved vascular maturity and reduced fibrotic response. These findings elucidate how the MWs of silk proteins shape angiogenic behavior and highlight the importance of MW tailoring for optimized tissue engineering applications.

## 1. Introduction

Engineered biomaterials have been widely used for various tissue regeneration, including skin, bone, cartilage, muscle, and so on ^[1]^. Angiogenesis is a fundamental process in tissue repair and wound healing, as newly formed blood vessels supply nutrients and oxygen while removing metabolic waste, thereby creating a favorable microenvironment for regeneration. In recent years, polymer-based biomaterials incorporating angiogenic cues—such as growth factors, therapeutic metal ions, and other pro-angiogenic agents—have demonstrated the ability to promote tissue vascularization ^[2]^. Alternatively, there is an increasing demand for bioactive polymers that possess inherent pro-angiogenic properties. To date, only a few naturally derived polymers have shown endogenous angiogenic activity ^[3]^, including collagen ^[4]^, gelatin ^[5]^, and hyaluronic acid ^[6]^.

Silk proteins derived from the cocoons of *Bombyx mori*, including silk sericin (SS) and silk fibroin (SF), have been widely utilized in tissue engineering in various forms over the past decade ^[7]^. In addition to their excellent processability, biocompatibility, and biodegradability, silk proteins have been shown to stimulate angiogenesis even in the absence of exogenous pro-angiogenic factors. For SS, multiple studies have demonstrated its ability to promote angiogenesis in murine models ^[8]^. Mechanistically, SS was reported to enhance vascular endothelial growth factor (VEGF) levels via MEK–ERK signaling in fibroblasts ^[8c]^, and to increase VEGF production through the HIF pathway in macrophages ^[8d]^. Regarding SF, there have been many reports showing its pro-angiogenic capacity in vivo ^[8b, 9]^. Mechanistically, SF was found to upregulated VEGF expression by activating NF-κB signaling in fibroblasts ^[9c]^.

Typically, a silk fibril in the cocoon consists of an outer layer of SS and an inner layer of SF. SS is a mixture of sericin proteins with molecular weights (MWs) ranging from 70 kDa to 400 kDa ^[10]^, whereas SF is a structural protein composed of a heavy chain (approximately 391 kDa) and a light chain (approximately 25 kDa) ^[11]^. A critical step in preparing SS- or SF-based biomaterials is isolating these proteins from the cocoons, which is commonly performed using chemical or heat treatments. However, such treatments inevitably reduce the MWs of the extracted SS ^[12]^ and SF ^[13]^. Importantly, changes in the MWs of silk proteins affect not only their physical properties ^[12b, 12c, 13d, 13e, 14]^, but also play a crucial role in determining their biological activities ^[15]^. Consequently, biomaterials fabricated from SS or SF with different MWs may exhibit distinct in vivo performance ^[12a, 15b, 16]^. Therefore, selecting silk proteins with specific MW distributions as building blocks becomes essential. Nevertheless, the relationship between the MWs of silk proteins and their angiogenic bioactivity remains largely unexplored.

In this study, to elucidate the impact of MWs on the angiogenic activities of *Bombyx mori* silk proteins, we employed alkaline hydrolysis to modulate their MW distributions and investigated both their direct and indirect effects on angiogenesis. Using endothelial cell tube formation assays and in vivo subcutaneous implantation experiments, we demonstrate that high-MW sericin (SSH) induces angiogenesis primarily through macrophage-mediated mechanisms involving mixed M1/M2-like phenotypes. In contrast, low-MW sericin (SSL) promotes angiogenesis by directly stimulating endothelial cells and by eliciting M2-like macrophages. For fibroin, the angiogenic activity of high-MW fibroin (SFH) is mainly associated with direct effects on endothelial cells rather than macrophages, whereas low-MW fibroin (SFL) enhances angiogenesis through both endothelial cell activation and M2-like macrophage involvement (**Scheme 1**).

**Scheme 1.**
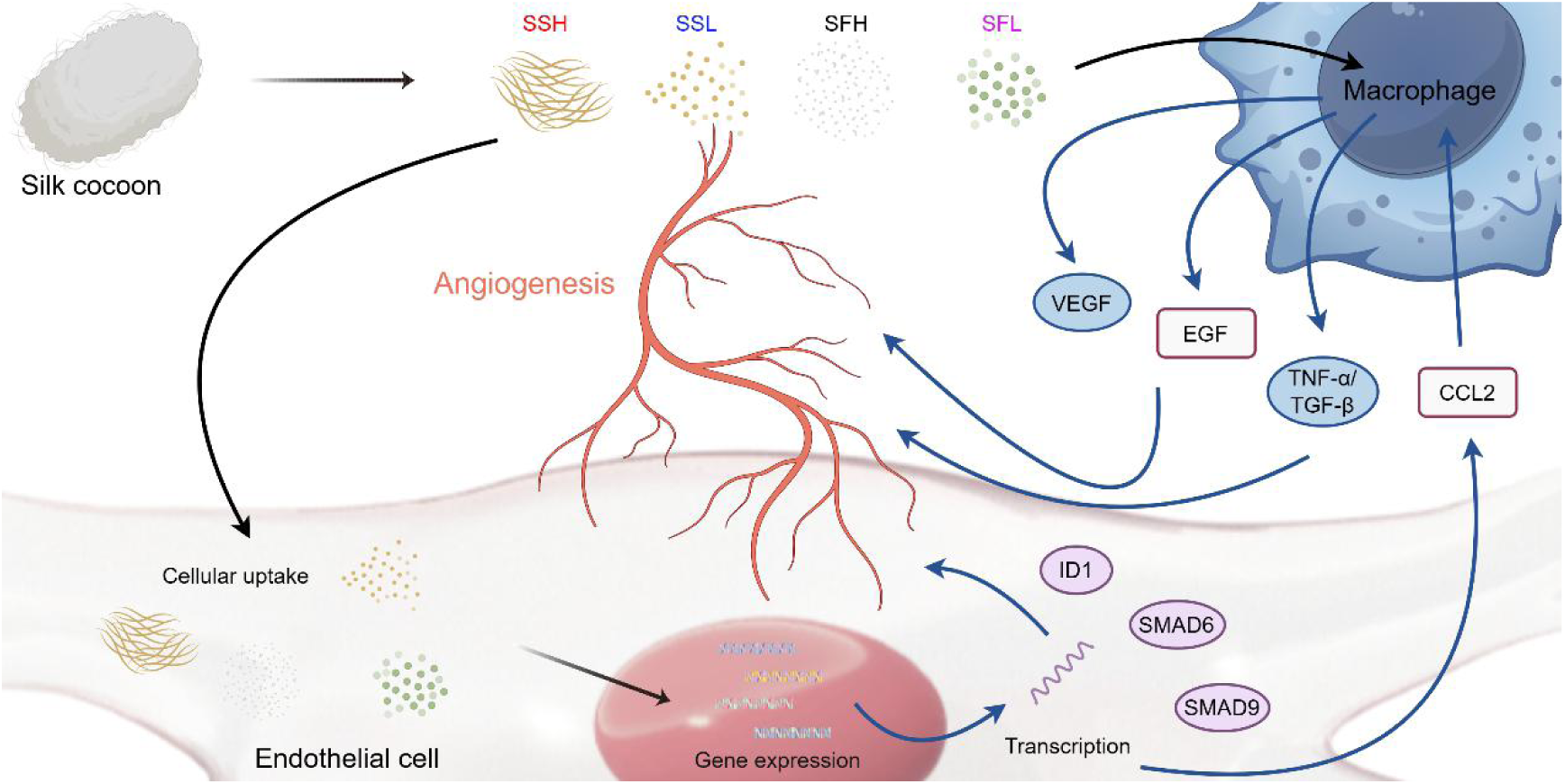
Preparation of silk sericin and silk fibroin with high and low-molecular weights, and their direct and indirect angiogenic effects. The scheme was created by Figdraw.

## 2. Materials and methods

### 2.1 Preparation of silk proteins with different MWs

To prepare high–molecular-weight silk sericin (SSH), *B. mori* silkworm cocoons (3.0 g) (XinDa Silk Co., Ltd., Guangdong, China) were cut into small pieces and boiled in deionized water for 30 min to extract SS. The remaining cocoon material were air-dried to calculate the extraction yield of SS (17.1%). The collected SS solution was centrifuged to remove insoluble aggregates and subsequently lyophilized, yielding SSH powder with an overall recovery of 83.9%. For the preparation of low–molecular-weight sericin (SSL), the freshly extracted SS solution was further boiled for an additional 60 min in 0.02 M Na₂CO₃. After hydrolysis, the solution was centrifuged and dialyzed against deionized water to remove salts. The resulting solution was freeze-dried to obtain SSL powder, with a final yield of 81.7%.

To prepare high–molecular-weight silk fibroin (SFH), *B. mori* silkworm cocoons (3.0 g) were cut into small pieces and boiled in 0.02 M Na₂CO₃ solution (Energy Chemical, Shanghai, China) for 30 min, followed by thorough rinsing with deionized water. This degumming process was repeated once to ensure complete removal of sericin. The remaining fibroin fibers were air-dried (yield: 66.3%) and subsequently dissolved in 9.3 M LiBr solution (Energy Chemical, Shanghai, China) at 60 °C for 4 h. The resulting SF solution was dialyzed against deionized water using a dialysis cassette (MWCO 3500 Da) for 3 days and centrifuged to remove insoluble aggregates. The clear solution was then freeze-dried to obtain SFH, with a final yield of 80.7%. To prepare low–molecular-weight fibroin (SFL), the SFH solution was further boiled in 0.02 M Na₂CO₃ for 60 min. As with SSL, the hydrolyzed solution was centrifuged, dialyzed against deionized water, and lyophilized to obtain SFL powder, yielding 73.0%.

### 2.2 Characterization of silk proteins

The molecular weight distributions of silk proteins were analyzed using sodium dodecyl sulfate–polyacrylamide gel electrophoresis (SDS-PAGE). Briefly, silk protein samples were mixed with an equal volume of loading buffer and boiled at 95 °C for 5 min before electrophoresis. After separation, the gels were stained using a fast silver staining kit (Solarbio, Beijing, China). The microscopic morphology of the silk proteins was examined by transmission electron microscopy (TEM, Talos L120C, Thermo Fisher Scientific). To evaluate protein solubility, increasing amounts of silk proteins were added to phosphate-buffered saline (PBS), vortexed, and centrifuged to collect the supernatants. Protein concentrations in the supernatants were quantified using a BCA protein assay kit (Beyotime, Shanghai, China). The size distribution of silk protein assemblies was determined by dynamic light scattering (DLS, Zetasizer Nano ZS90, Malvern, UK). Fourier transform infrared spectroscopy (FT-IR, VERTEX 70, Bruker, USA) was performed to analyze functional groups and secondary structures, with spectra collected in the range of 4000–400 cm⁻¹. Circular dichroism (CD, Chirascan V100, Applied Photophysics, UK) was further employed to assess the molecular conformation of silk proteins in solution.

### 2.3 Cell culture

Human umbilical vein endothelial cells (HUVECs) were cultured in an endothelial cell complete medium (Zhongqiao Xinzhou Biotechnology Co., Ltd, Shanghai, China). RAW 264.7 cells were maintained in high-glucose DMEM (Gibco, USA) supplemented with 10% fetal bovine serum (FBS) and 1% penicillin/streptomycin. All cells were cultured at 37°C in a humidified atmosphere containing 5% CO_2_. For conditioned medium collection, RAW 264.7 cells were treated with or without 0.01% (w/v) silk proteins in the medium. After incubation for 24 h, the culture supernatant was collected and centrifuged to remove dead cells and debris before use.

### 2.4 Cell counting kit-8 (CCK-8) assay

Cell proliferation was assessed using the CCK-8 kit (Beyotime, Shanghai, China). Cells were seeded into 96-well plates at a density of 5×10^3^ cells per well. All types of cells were treated with a gradient of silk protein concentrations. At the designated time points, the medium was replaced with a 10% (v/v) CCK-8 solution. After incubation at 37 °C for 1 h, absorbance was measured at 450 nm using a microplate reader (Multiskan FC, Thermo, USA).

### 2.5 Live/dead cell staining

Cell viability was assessed using the Calcein/PI cell viability and cytotoxicity assay kit (Beyotime, Shanghai, China). Cells were seeded into 96-well plates at a density of 5×10^3^ cells per well. Fluorescence images were captured using a microscope (TS100, Nikon, Japan), where live cells fluoresced green and dead cells fluoresced red.

### 2.6 Cell migration

Cell migration was assessed using a scratch assay. HUVECs were seeded in 24-well plates and cultured to ∼90% confluence. A scratch was made in the center of the cell monolayer using a 1-mL pipette tip. Cells were then incubated with growth medium containing 0.1% (w/v) silk proteins or without silk proteins as a control. Images were captured at 0 and 24 h after staining with Calcein AM (Beyotime, Shanghai, China) using a fluorescence microscope (Nikon, Japan), and migration was quantified using ImageJ software.

### 2.7 Tube formation assay

A tube formation assay was performed to assess the angiogenic potential of HUVECs. Pre-cooled 96-well plates were coated with Matrigel (356230, Corning, USA) and allowed to solidify at 37 °C for 3 h. Subsequently, 3 × 10⁴ HUVECs were seeded into each well and incubated with growth medium containing 0.1% (w/v) silk proteins or various conditioned medium from macrophages for 6 h. Cells were then stained with Calcein AM (Beyotime, Shanghai, China), and images were captured using a fluorescence microscope (TS100, Nikon, Japan). The number of nodes and total tube length were quantified using the angiogenesis analyzer plugin in ImageJ software.

### 2.8 Cellular uptake

To examine the cellular uptake of silk proteins, they were labeled with fluorescein isothiocyanate (FITC). Briefly, 50 mg of silk proteins were dispersed in 20 mL of deionized water, and the pH was adjusted to 8.2. FITC solution was then added, and the mixture was stirred in the dark for 12 h. The resulting solution was dialyzed against deionized water for 3 days using a dialysis cassette to remove unreacted FITC. RAW 264.7 cells were incubated for 24 h with medium containing 0.01% (w/v) FITC-labeled silk proteins. After incubation, cells were stained with DAPI (Beyotime, Shanghai, China) and imaged using a confocal laser scanning microscope (CLSM, Leica, Germany).

### 2.9 RT-qPCR

To assess cytokine gene expression, real-time quantitative polymerase chain reaction (RT-qPCR) was performed. Cells were incubated for 24 h in a growth medium containing either 0.1% (w/v) silk proteins (HUVECs) or 0.01% (w/v) silk proteins (RAW 264.7 cells) before total RNA extraction. Reverse transcription was performed using PrimeScript™ RT Master Mix (RR047A). The primers used in this study are listed in Table S1.

### 2.10 Enzyme-linked immunosorbent assay (ELISA)

Cells were incubated with a growth medium containing 0.1% (w/v) silk proteins (HUVECs) or 0.01% (w/v) silk proteins (RAW 264.7 cells) for 24 h. As a positive control, RAW 264.7 cells were treated with 1 μg/mL LPS (Beyotime, Shanghai, China). After incubation, the culture medium was collected and centrifuged to remove dead cells and debris. The resulting supernatant was stored at 4 °C until analysis. Cytokine levels, including TNF-α, TGF-β, and VEGFA, were quantified according to the manufacturer’s protocols (Elabscience, China).

### 2.11 RNA sequencing

HUVECs incubated with a growth medium containing 0.1% (w/v) silk proteins for 24 h were collected for RNA sequencing. RNA sequencing was performed by Kidio Biotechnology Co., Ltd (Guangzhou, China). All data analysis was performed by Omicsmart cloud platform (www.omicsmart.com).

### 2.12 Western blot

RAW 264.7 cells were incubated with a growth medium containing 0.01% (w/v) silk proteins for Western blotting. As a positive control, cells treated with LPS (1 μg/mL) (Beyotime, Shanghai, China) were used. Total protein was extracted from the cells, and its concentration was measured using a BCA protein assay kit (Beyotime, Shanghai, China). The proteins were separated via SDS-PAGE and transferred to a polyvinylidene difluoride (PVDF) membrane. The membrane was incubated overnight at 4 °C with primary antibodies against iNOS (80517-1-RR, Proteintech, Wuhan, China), Arg1 (AF1381, Beyotime, Shanghai, China), VEGF (AF1309, Beyotime, Shanghai, China), and α-tubulin (AF2827, Beyotime, Shanghai, China). Subsequently, the membrane was incubated with a secondary antibody (A0208, Beyotime, Shanghai, China) and visualized using an electrochemiluminescence detection reagents.

### 2.13 Immunofluorescence assay

To visualize the expression of polarization markers of RAW 264.7 cells, the cells (1.5 × 10^5^) seeded in confocal dishes were incubated with silk proteins (0.01% (w/v)) or LPS (1 μg/mL). After 24 h incubation, the cells were washed with PBS and fixed in 4% paraformaldehyde for 10 min, permeabilized with 0.1% (v/v) Triton X-100 for 5 min and blocked with 1% (w/v) BSA for 30 min. Following washing with PBS, the cells were incubated overnight at 4 ℃ with the primary antibody against CD86 (83213-1-RR, Proteintech, Wuhan, China) and CD206 (81525-1-RR, Proteintech, Wuhan, China). Then, the cells were incubated with Alexa Fluor™ 488-conjugated secondary antibody at room temperature and stained with DAPI before imaging by CLSM. Quantitative analysis was performed using ImageJ software.

### 2.14 Limulus amoebocyte lysate (LAL) endotoxin assay

Endotoxin content of the four silk proteins was measured using a chromogenic LAL endotoxin assay kit (Beyotime, Shanghai, China), and the absorbance was read at a wavelength of 545 nm using a microplate reader (Multiskan FC, ThermoFisher, USA). Sample detection concentration was 0.01% (w/v).

### 2.15 In vivo subcutaneous injection mice model

All experimental procedures were approved by South China Agricultural University and conducted in compliance with the guidelines for the care and use of laboratory animals (Application number: 2024B183). Eight-week-old female C57BL/6 mice were obtained from Bestest Biotechnology Co., Ltd. (Zhuhai, China). SSH and SSL were injected into the mice (n=6). PBS was used as control. Before the procedure, anesthesia was administered. Using a syringe, 200 μL of sericin solution at a concentration of 0.5% (w/v) was injected subcutaneously. The mice were monitored until they recovered from anesthesia and were then housed for 7 and 14 days. At each time point, the mice were euthanized, and the tissues surrounding the injection site were excised and collected. The collected samples were fixed in 4% paraformaldehyde overnight.

### 2.16 Histological analysis, immunofluorescence and immunochemistry

After the samples were fixed in 4% paraformaldehyde and embedded in paraffin wax, sections of each sample (3-5 μm thick) were prepared and mounted onto slides. Angiogenesis on the injection surface was assessed by immunofluorescence staining of the vascular marker CD31 (80530-1-RR, Proteintech, Wuhan, China) and by immunohistochemistry staining of VEGF (AF1309, Beyotime, Shanghai, China) and α-SMA (GB111364, Servicebio, Wuhan, China). The sections were stained with H&E to evaluate the fibrotic capsule and inflammatory cell infiltration. Macrophage infiltration at the injection site was detected via immunohistochemistry staining of the macrophage marker F4/80 (GB113373, Servicebio, Wuhan, China). M1 and M2-like macrophages were identified by immunofluorescence staining using antibodies against CD86 (83213-1-RR, Proteintech, Wuhan, China) and CD206 (81525-1-RR, Proteintech, Wuhan, China), respectively. H&E images and immunohistochemistry images were captured using a light microscopy, while immunofluorescence images were obtained using fluorescence microscopy. Quantitative analysis was performed using ImageJ software.

### 2.17 Data analysis

Data were expressed as the mean ± standard deviation (SD). Statistical comparisons and nonlinear regressions were performed using Prism 9.0 (GraphPad Software, Inc.). Data normality was analyzed by Shapiro–Wilk test. The significance of differences was assessed using Student’s t-test or one-way analysis of variance (ANOVA), with a *P*-value less than 0.05 considered statistically significant.

## 3. Results

### 3.1 Characterization of silk proteins with different molecular weights

First, silk proteins with distinct MWs were prepared by controlling the extent of alkaline hydrolysis. The MW distributions of SS and SF were determined by sodium dodecyl-sulfate polyacrylamide gel electrophoresis (SDS-PAGE). As shown in **Figure 1a**, SSH and SFH exhibited a smeared band ranging from ∼40 kDa to ∼180 kDa, whereas SSL and SFL displayed bands less than ∼25 kDa, respectively, indicating that alkaline hydrolysis is an effective method to tune the MWs of silk proteins. Solubility of the four silk proteins in PBS was next evaluated. As shown in Figure S1, SS and SF displayed distinct solubility profile. The dissolution rate of SSH sharply decreased from 30% to 5% as concentration increased at room temperature, whereas SSL fully dissolved in PBS up to 0.5% (w/v) and reached ∼75% dissolution at 1% (w/v). Both SFH and SFL fully dissolved even at 1% (w/v), demonstrating superior solubility compared with SS.

**Figure 1.**
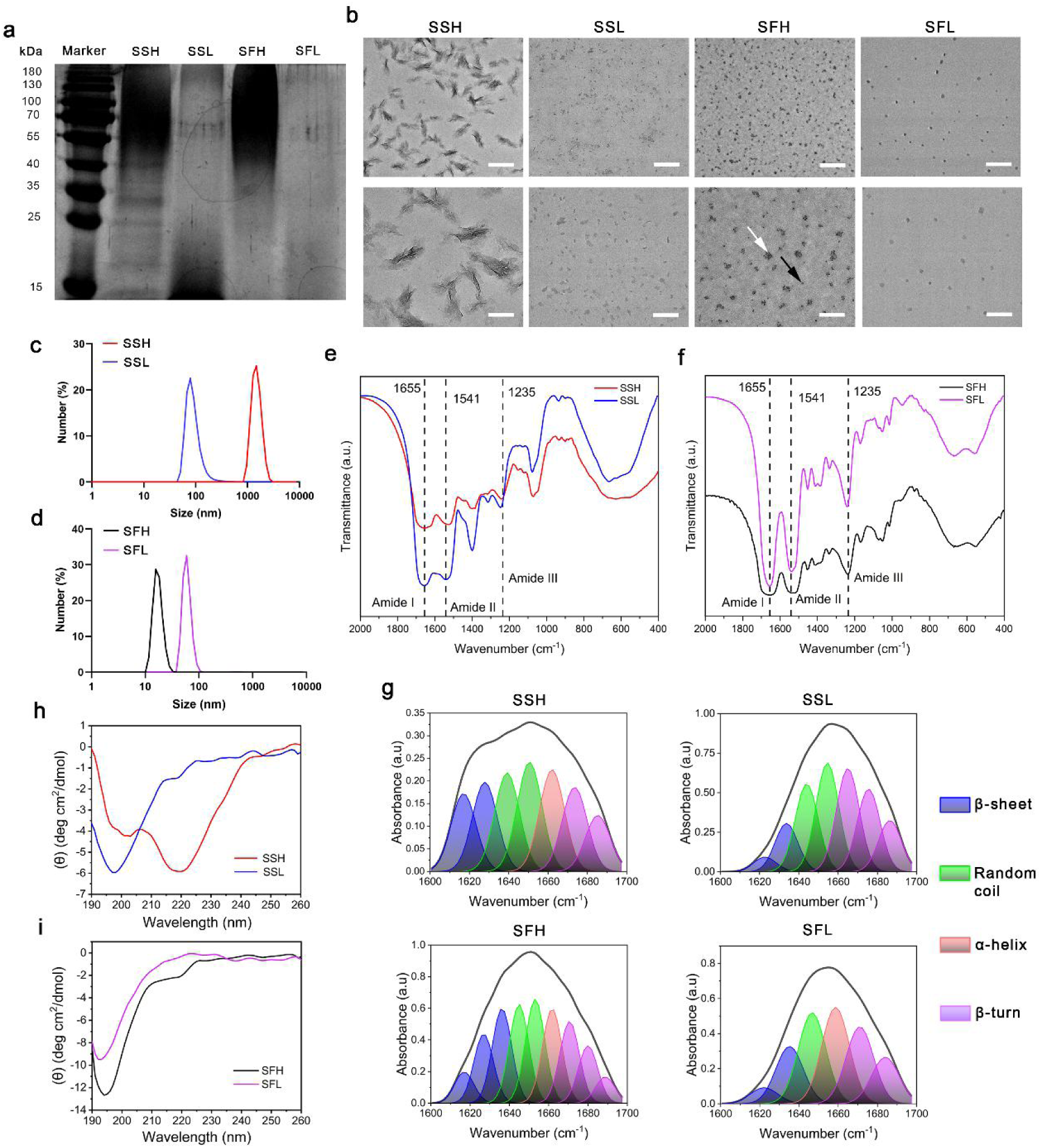
Characterization of silk proteins with different molecular weights. (a) SDS-PAGE images of the four silk proteins. (b) TEM images of the four silk proteins. Black arrow represents single particle, and white arrow represents aggregated particles. Scale bar, 500 nm for the upper panel and 200 nm for the lower panel. (c, d) Size distributions of silk sericin (SSH and SSL) (c) and silk fibroin (SFH and SFL) (d) in PBS measured by dynamic light scattering (DLS). (e, f) FTIR spectra of silk sericin (c) and silk fibroin (d). (g) Deconvolution of FTIR spectra in (e, f). (h, i) Circular dichroism (CD) spectra of silk sericin (SSH and SSL) (h) and silk fibroin (SFH and SFL) (i) in PBS.

The morphology of silk proteins was further examined by transmission electron microscopy (TEM). Notably, SSH and SSL displayed pronounced structural differences: SSH formed bundled nanofibers several hundred nanometers in length, while SSL appeared as irregular nanoparticles smaller than 100 nm **(Figure 1b)**. For SF, SFH formed nanoaggregates composed of smaller particles, whereas SFL appeared as unstructured nanoparticles (Figure 1b). Dynamic light scattering (DLS) analysis confirmed these observations: SSH had an average size of ∼2 μm, SSL ∼70 nm, SFH ∼20 nm, and SFL ∼60 nm (Figure 1c). Collectively, these results suggest that MW alterations exert a more pronounced structural effect on SS than on SF.

Given the structural differences induced by MW reduction, we analyzed the conformational changes of silk proteins using Fourier-transform infrared spectroscopy (FTIR) and circular dichroism (CD). The infrared spectral region between 1700–1200 cm⁻¹ corresponds to peptide backbone absorption: amide I (1700–1600 cm⁻¹), amide II (1600–1500 cm⁻¹), and amide III (1350–1200 cm⁻¹), commonly used to assess protein secondary structures. As shown in **Figure 1e**, the amide I band of SSL peaked at 1655 cm^-1^, corresponding to conformation of random coil. In contrast, SSH showed a primary peak at 1655 cm⁻¹ accompanied by a shoulder at 1625 cm⁻¹, reflecting a mixture of β-sheet and random coil structures. For SF, both SFH and SFL displayed amide I peaks at 1655 cm⁻¹, corresponding predominantly to random coil (Figure 1f). To quantify secondary structure content, the amide I bands were deconvoluted, revealing a clear transition from β-sheet to random coil/helix as MW decreased (Figure 1g and Figure S2). CD analysis of silk proteins in solution further confirmed these results: SSH exhibited a strong negative band near 218 nm, consistent with β-sheet conformation, whereas SSL showed a negative band around 198 nm, characteristic of random coil (Figure 1h). Both SFH and SFL presented dominant negative bands near 195 nm, indicating random coil as the prevailing structure, although SFH displayed a minor β-sheet signal near 218 nm (Figure 1i). Collectively, these results suggest that MW reduction markedly disrupts the β-sheet content and fiber-forming capacity of SS, while exerting a comparatively mild effect on the secondary structure of SF.

### 3.2 Direct angiogenic effect of silk proteins with different molecular weights on HUVECs

To explore the angiogenic potential of silk proteins, we investigated their effects on key cellular behaviors of HUVECs, including viability, migration, and tube formation. First, HUVEC proliferation was assessed using the cell counting kit-8 (CCK-8) assay after exposure to silk protein concentrations ranging from 0.01 to 0.2% (w/v) for 24 h (Figure 2a). The results indicated no significant cytotoxicity at any concentration. However, there was a significant decrease in cell proliferation rate following treatments of SSL (*P*<0.01) and SFH (*P*<0.05) via ANOVA trend analysis. Considering the proliferation rate of HUVECs incubated by the four silk proteins at 0.1% (w/v) was above 90%, 0.1% (w/v) was selected as the working concentration for subsequent experiments (Figure 2b).

**Figure 2.**
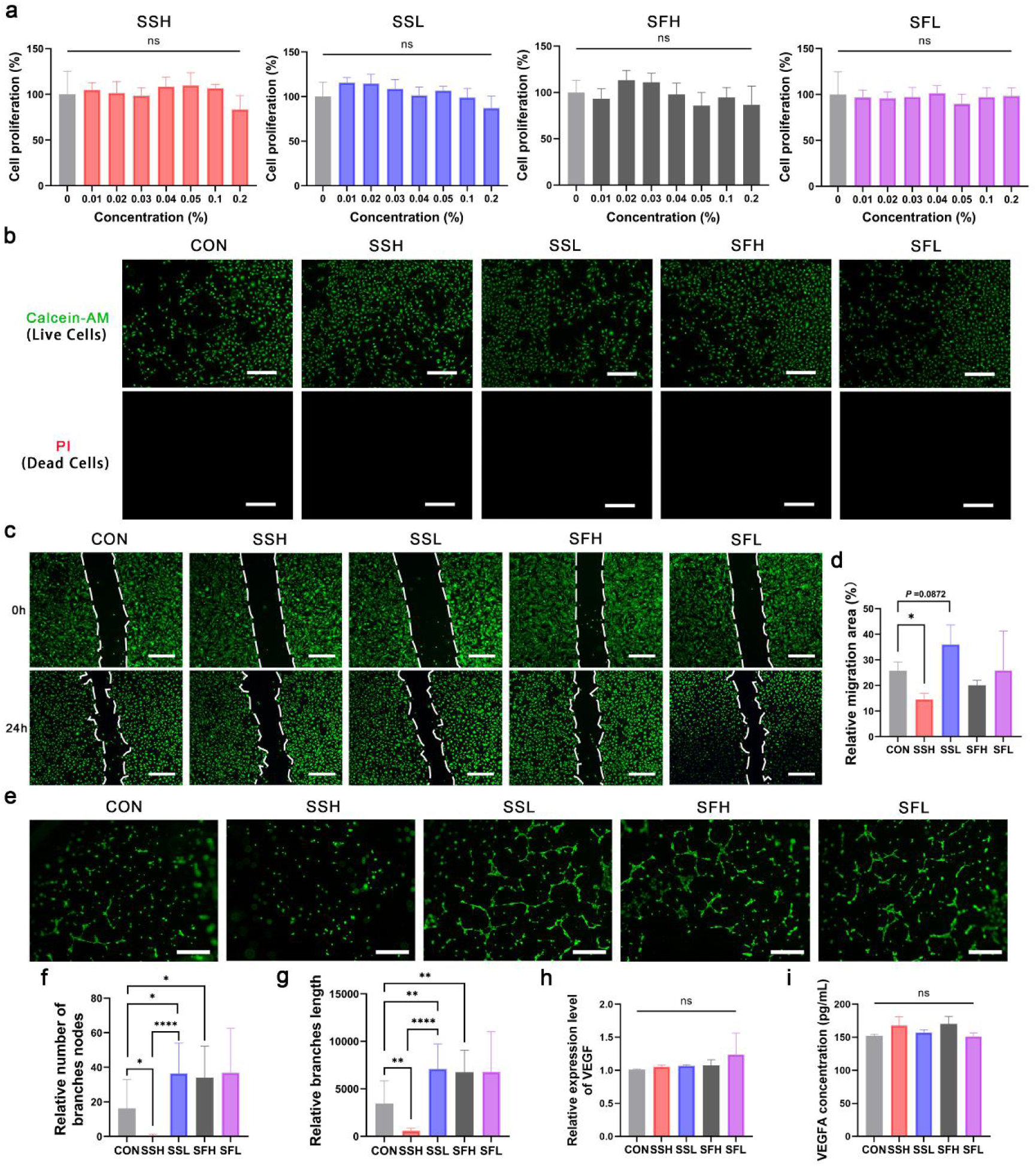
Silk proteins with different molecular weights have distinct effects on the angiogenesis of HUVECs in vitro. (a) Cell proliferation profile of HUVECs treated by 0.01 to 0.2% (w/v) silk proteins for 24 h (n=5). (b) Calcium-AM staining images of live cells (green) and PI staining of dead cells (red) in HUVECs treated with 0.1% (w/v) silk proteins for 24 h (n=3). (c, d) Scratch assay of HUVECs and quantitative analysis for cellular migration at 0 and 24 h. Cells were treated with growth medium containing 0.1% (w/v) silk proteins or with growth medium alone as a control. (e-g) Representative images and quantitative analysis of tube formation of HUVECs. The cells were treated with growth medium containing 0.1% (w/v) silk proteins or with growth medium alone as a control for 6 h. (f) Relative number of branch nodes (n=3). (g) Relative branch length (n=3). (h, i) RT-qPCR analysis of VEGF (h) and ELISA analysis of VEGFA (i) following incubation of HUVECs with 0.1% (w/v) silk proteins for 24 h (n=3). Scale bar, 50 μm. Data represent mean ± SD, \**P* < 0.05, \*\**P* < 0.01, \*\*\*\**P* < 0.0001, ns: not significant.

Next, a scratch assay was performed to assess HUVEC migration (Figure 2c, d). Interestingly, SSH inhibited cell migration compared to the control group (CON) (*P*<0.05), whereas SSL slightly, though not significantly (*P*=0.0872), enhanced HUVEC migration. Both SFH and SFL had minimal effects on migration. To further evaluate the angiogenic potential of silk proteins, a tube formation assay was conducted. Capillary-like structures were more pronounced in the presence of SSL, SFH, and SFL compared to CON (Figure 2e). Quantitative analysis of node number and tube length confirmed that SSL, SFH, and SFL promoted angiogenesis, while SSH exhibited the opposite effect (Figure 2f, g). Considering VEGF as a key angiogenic factor, its expression in HUVECs was examined by RT-qPCR and ELISA, revealing no significant differences among the four silk proteins (Figure 2h, i). These results highlight the positive roles of SSL, SFH, and SFL, and the inhibitory effect of SSH on HUVEC tube formation.

### 3.3 Silk proteins with different molecular weights trigger distinct transcriptional pathways in HUVECs

To gain deeper insights, we conducted RNA sequencing to analyze the total mRNA expression of HUVECs following silk protein treatments. The principal component analysis (PCA) results showed clear clustering patterns among the different groups (Figure 3a). Notably, SSH and SSL were distinctly separated, indicating substantial differences in their overall gene expression profiles, whereas SFH and SFL clustered closely, suggesting relatively minor variations between them. As revealed in Figure 3b, a total of 70 genes were significantly differentially expressed in SSH-treated cells compared to CON, with up- and down-regulated genes being roughly equal in number. SSL induced more upregulated genes relative to CON, a trend also apparent when directly comparing SSH with SSL. In contrast, SFH and SFL exhibited minimal differential gene expression, a pattern that persisted when comparing SFH with SFL. These results indicate that silk proteins elicit a relatively mild transcriptional response in HUVECs overall, with SS provoking a greater effect than SF.

**Figure 3.**
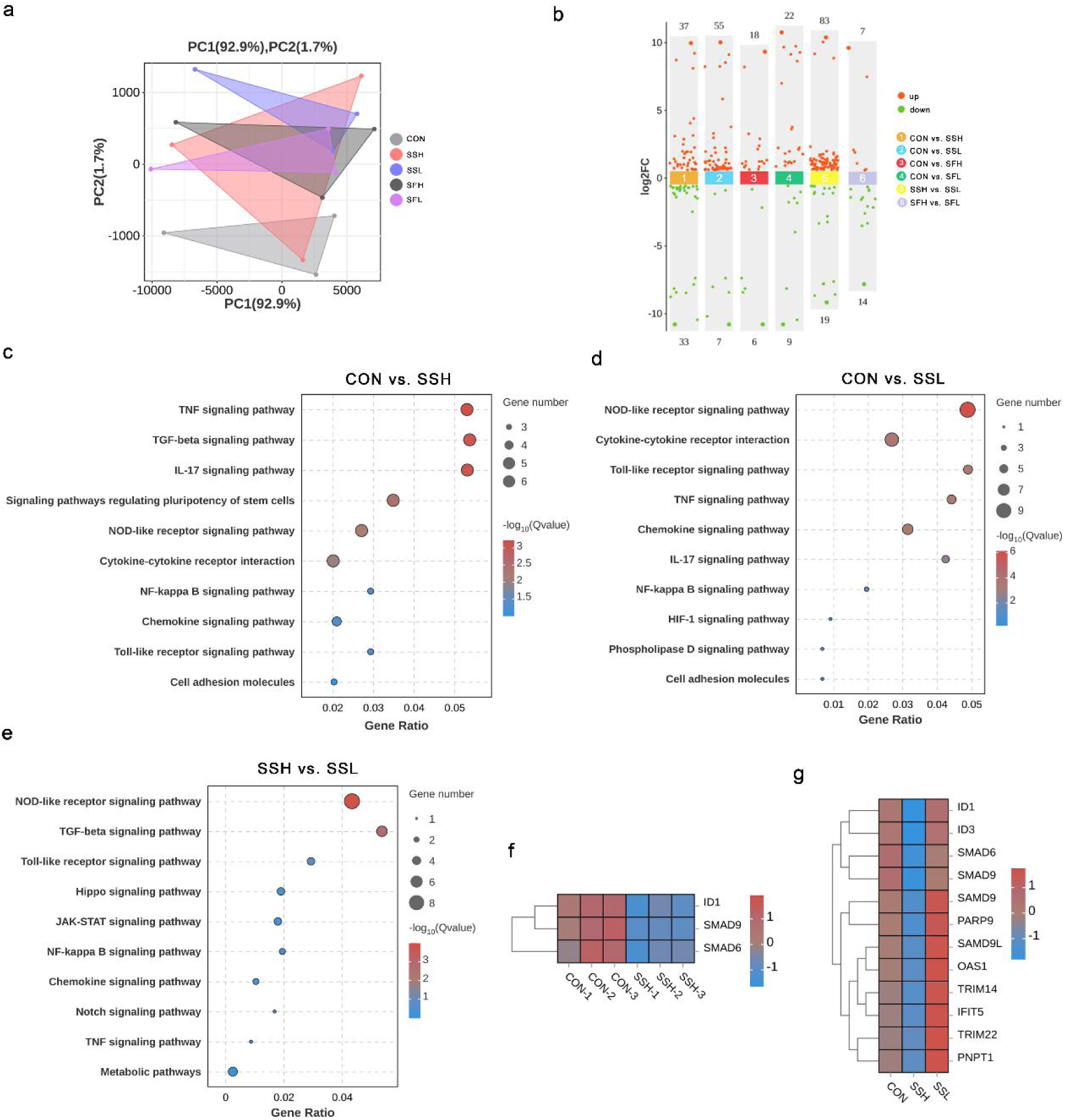
Transcriptomic analysis of HUVECs treated with the four silk proteins with different molecular weights. (a) Sample differences by principal component analysis (PCA). (b) Scatter plot of different genes in multiple groups. (c-e) KEGG analysis of different signaling pathways in SSH and SSL-treated cells. (f, g) The heatmap of differential genes involved in SSH and SSL-treated cells. n=3.

Given that SS induced more differentially expressed genes, we focused the main transcriptome analysis on SSH and SSL. Kyoto Encyclopedia of Genes and Genomes (KEGG) pathway analysis revealed that, compared to CON, both SSH and SSL significantly activated inflammation-related pathways, including TNF, TGF-β, IL-17, and NOD-like receptor signaling (Figure 3c, d). Compared with SSH, SSL specifically activated key pathways such as NOD-like receptor and TGF-β signaling (Figure 3e). We further screened differentially expressed genes using a false discovery rate (FDR) cutoff. Notably, *Id1*, *Smad6*, and *Smad9*—genes associated with angiogenesis—were downregulated in SSH-treated HUVECs relative to CON (FDR < 0.05; Figure 3f). It implies that the genes may be responsible for the impaired tube formation of SSH-treated HUVECs. When comparing SSH and SSL with an FDR threshold of 0.1, SSH and SSL displayed opposing effects on HUVECs, not only due to downregulation of *Id1*, *Smad6*, and *Smad9* by SSH, but also because SSL significantly upregulated angiogenesis-related genes such as *Samd9* and *Samd9l* (Figure 3g). Similarly, KEGG analysis of SFH and SFL revealed minimal gene expression differences (Figure S3-5). Collectively, the differential regulation of key genes, including *Id1* and *Smad6/9*, likely underlies the contrasting angiogenic effects of SSH and SSL on HUVECs.

### 3.4 Indirect effect of silk proteins with different molecular weights on angiogenesis via macrophages

Macrophages are key players in tissue injury, repair, and regeneration. Along with endothelial cells, they represent two crucial cell types that play distinct yet interconnected roles in maintaining tissue homeostasis. Given that macrophages are essential regulators of angiogenesis during tissue repair, we next investigated whether they contribute to silk protein-mediated angiogenesis. First, we assessed the proliferation of RAW 264.7 cells treated with silk proteins. No significant stimulatory or inhibitory effects were observed within the tested concentration range (Figure 4a). Cellular uptake of the four silk proteins was then examined by CLSM (Figure S6), showing that low-MW silk proteins were more readily endocytosed by macrophages. To evaluate the influence of macrophage-derived factors on endothelial tube formation, RAW 264.7 cells were incubated with silk proteins, and the resulting conditioned medium were applied to HUVECs (Figure 4b). Unexpectedly, conditioned media from SSH, SSL, and SFL markedly promoted tube formation of HUVECs, as demonstrated by fluorescence microscope and quantitative analysis (Figure 4c-e).

**Figure 4.**
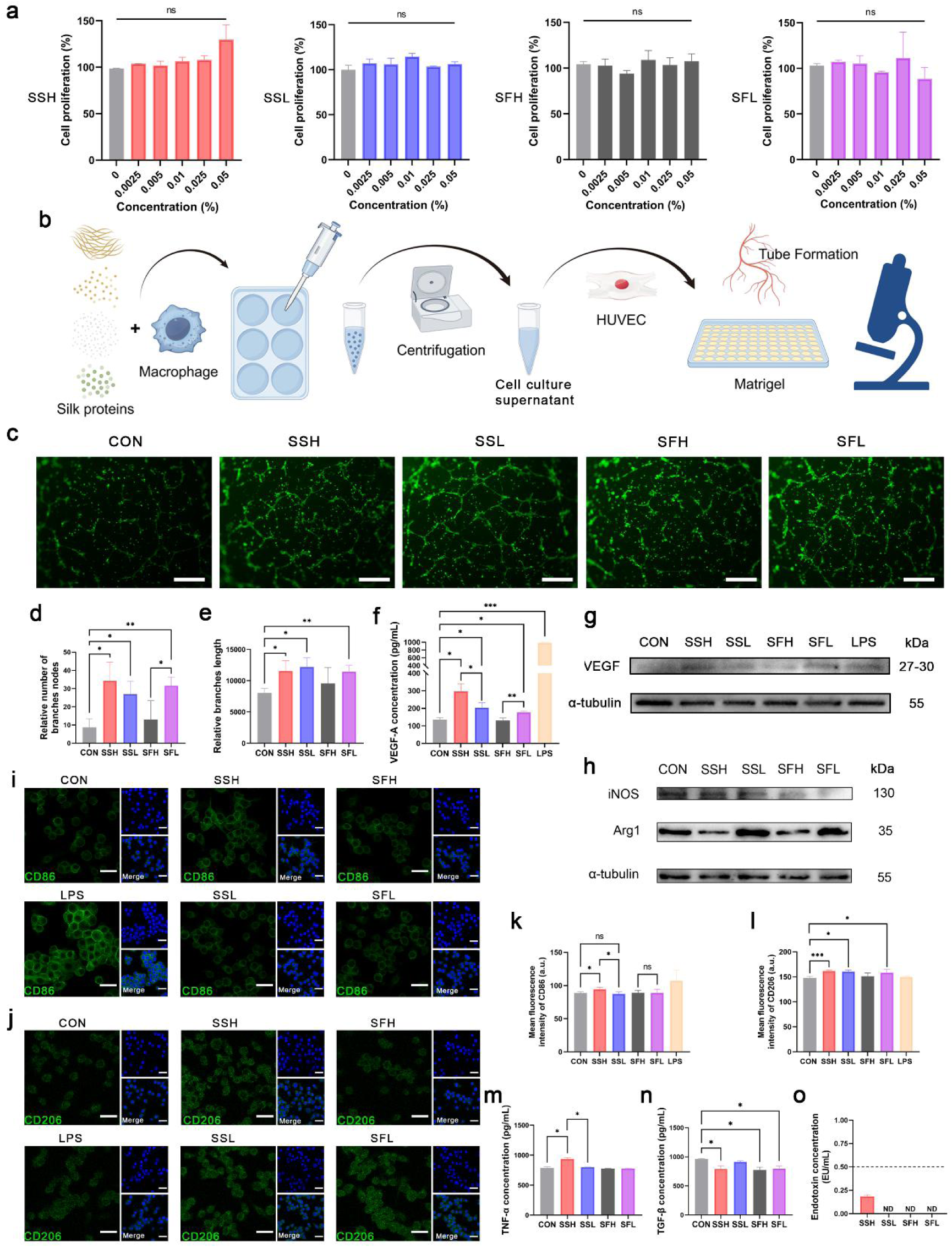
Effect of silk proteins with different MWs on macrophage-mediated angiogenesis. (a) Cell proliferation profile of RAW 264.7 cells incubated with 0.0025% to 0.05% (w/v) silk proteins for 24 h (n=5). (b) Scheme of the indirect tube formation assay, which was created by Figdraw. (c) Representative images of tubes formed by HUVECs treated with different conditioned media from silk protein-incubated RAW 264.7 cells for 6 h. Scale bar, 50 μm. (d, e) Quantitative analysis of tube forming of HUVECs measured by relative number of branch nodes (d) and relative branch length (e). (f) Concentration of VEGFA in RAW 264.7-derived conditioned media determined by ELLISA. (g) Western blot of VEGF expression in RAW 264.7 cells treated by the four silk proteins for 24 h. (h) Western blot of iNOS and Arg1 expression in RAW 264.7 cells treated by 0.01% (w/v) silk proteins for 24 h. (i-l) immunofluorescence images of CD86 (k) and CD206 (l) expression in RAW 264.7 cells incubated with 0.01% (w/v) silk proteins for 24 h, and quantitative analysis of fluorescence intensity of CD86 (m) and CD206 (n) (n=3). Scale bar, 30 μm. (m, n) Concentration of cytokines of TNF-α (i), and TGF-β (j) secreted by RAW 264.7 cells incubated with 0.01% (w/v) silk proteins for 24 h. (o) The content of endotoxin in silk proteins by limulus amoebocyte lysate (LAL) assay (n=3). Data represent mean ± SD, \**P* < 0.05, \*\**P* < 0.01, \*\*\**P* < 0.001, ns: not significant, ND, not detected.

To probe the underlying mechanism, we first measured VEGFA levels in the supernatant of silk-treated RAW 264.7 cells. Lipopolysaccharide (LPS) was used as a positive control due to its potent VEGF-inducing effect. Compared to CON, SSH, SSL, and SFL treatments significantly increased VEGFA secretion (*P*<0.05), with SSH exhibiting the highest concentration (Figure 4f). Consistently, western blot analysis confirmed elevated VEGF expression in RAW 264.7 cells treated with SSH, SSL, and SFL relative to CON (Figure 4g). Additionally, the mRNA level of another angiogenic factor, epidermal growth factor (EGF), was significantly increased upon treatment with SSH, SSL, and SFL (Figure S7). To examine the endothelial response, HUVECs were treated by conditioned medium from silk-treated RAW 264.7 cells, and the expression of C-C chemokine ligand 2 (CCL2), a chemoattractant for immune cells such as macrophages, was assessed. It is revealed that SSH-, SSL-, and SFL-derived conditioned medium markedly upregulated CCL2 expression in HUVECs compared to CON (*P*<0.01) (Figure S8).

To investigate whether the macrophage phenotype influences the VEGF secretion, we examined M1 and M2 markers in RAW 264.7 cells using western blot and immunofluorescence. Western blot analysis showed that SSH upregulated iNOS, an M1 marker, whereas SSL increased Arg1, an M2 marker. SFH downregulated Arg1 expression, while SFL upregulated it (Figure 4h). Consistently, immunofluorescence revealed that SSH induced increased signals for both CD86 (M1) and CD206 (M2), whereas SSL and SFL selectively enhanced CD206 fluorescence (Figure 4i-l). The level of TNF-α was higher in SSH-treated cells than in both CON and SSL groups (*P*<0.05) (Figure 4m), whereas TGF-β was decreased in SSH-treated cells (P<0.05) but remained largely unchanged in the SSL group (*P*>0.05) (Figure 4n). Given the M1-inducing effect of SSH, we measured endotoxin levels and detected trace amounts (below 0.2 EU/mL) only in SSH (Figure 4o). Collectively, these results suggest that SSH-mediated HUVEC tube formation may arise from M1/M2-like macrophage polarization, whereas SSL- and SFL-mediated angiogenesis is primarily associated with M2-like macrophage polarization.

### 3.5 Differential angiogenic effects of silk sericin with different molecular weight in a subcutaneous mouse model

Given the divergent effects of SSH and SSL on angiogenesis, we employed a subcutaneous implantation mouse model to evaluate the effect of MWs of SS on neovascularization. Skin tissues surrounding the injection sites were collected on 7 and 14 days post-implantation. Tissue vascularization was first assessed via immunofluorescence staining for CD31. Both SSH and SSL significantly enhanced angiogenesis on day 7, with SSH showing a more pronounced effect. By day 14, no significant differences in vessel density among SSH, SSL, and CON were observed, suggesting a return toward homeostasis (Figure 5a, b). To assess vascular stability and maturation, α-SMA staining was performed. SSL-treated tissues exhibited higher α-SMA expression than SSH on both days 7 and 14 (*P*<0.05), indicating that SSL promotes the formation of more mature vessels in vivo (Figure 5c, d). Consistent with vessel density, both SSH and SSL elevated VEGF levels on day 7 (*P*<0.05, *P*<0.01) (Figure 5e, f).

**Figure 5.**
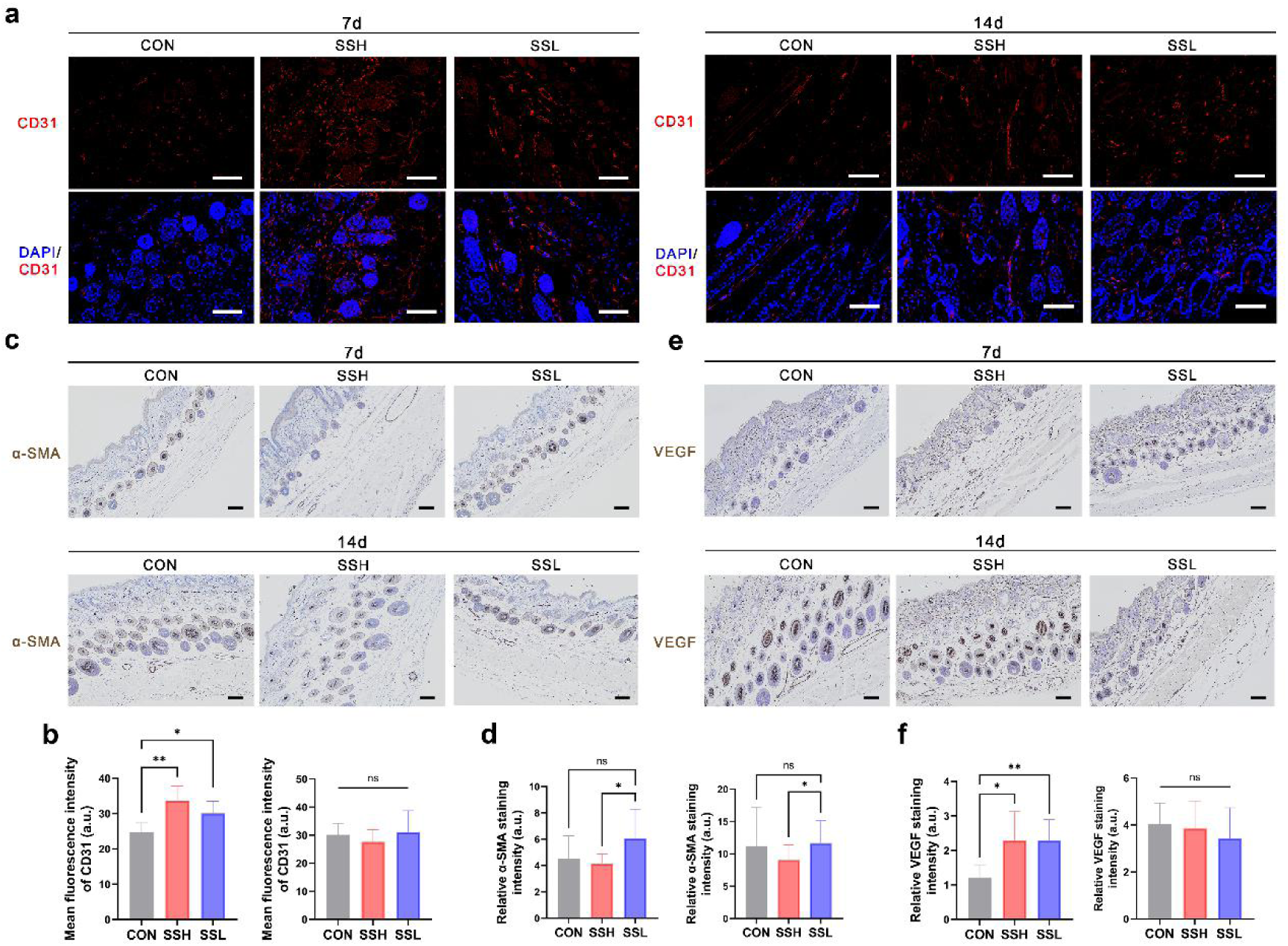
Pro-angiogenic effect of SSH and SSL in vivo. (a, b) Immunofluorescent staining and quantitative analysis of CD31 on 7 and 14 days. (c-f) Immunochemistry images and quantitative analysis by staining α-SMA (c, d) and VEGF (e, f) on 7 and 14 days. Scale bar, 100 μm. Data represent mean ± SD, n=6 mice per group, \**P* < 0.05, \*\**P* < 0.01, ns: not significant.

Next, we assessed the immune response and fibrosis surrounding the implantation sites. H&E staining revealed substantial infiltration of inflammatory cells and fibrin deposition around the SSH-treated sites on day 7, whereas SSL-treated sites exhibited a similar cell distribution to CON (Figure 6a, b). It suggests that SSL elicited a weaker immune response, possibly due to its improved hydrophilicity. Immunohistochemical analysis of F4/80-labeled macrophages showed a trend toward increased macrophage infiltration in both SSH and SSL groups, with SSH in the highest intensity, although differences were not statistically significant (Figure 6c, d). Immunofluorescence further characterized macrophage phenotypes around the implantation sites. While CD86 expression was comparable across all groups, SSL-treated tissues exhibited a significantly higher proportion of CD206+ macrophages compared to SSH (*P*<0.05) (Figure 6e-g). Overall, these in vivo results demonstrate that both SSH and SSL promote angiogenesis, with SSL more effectively enhancing vascular maturation and reducing fibrosis.

**Figure 6.**
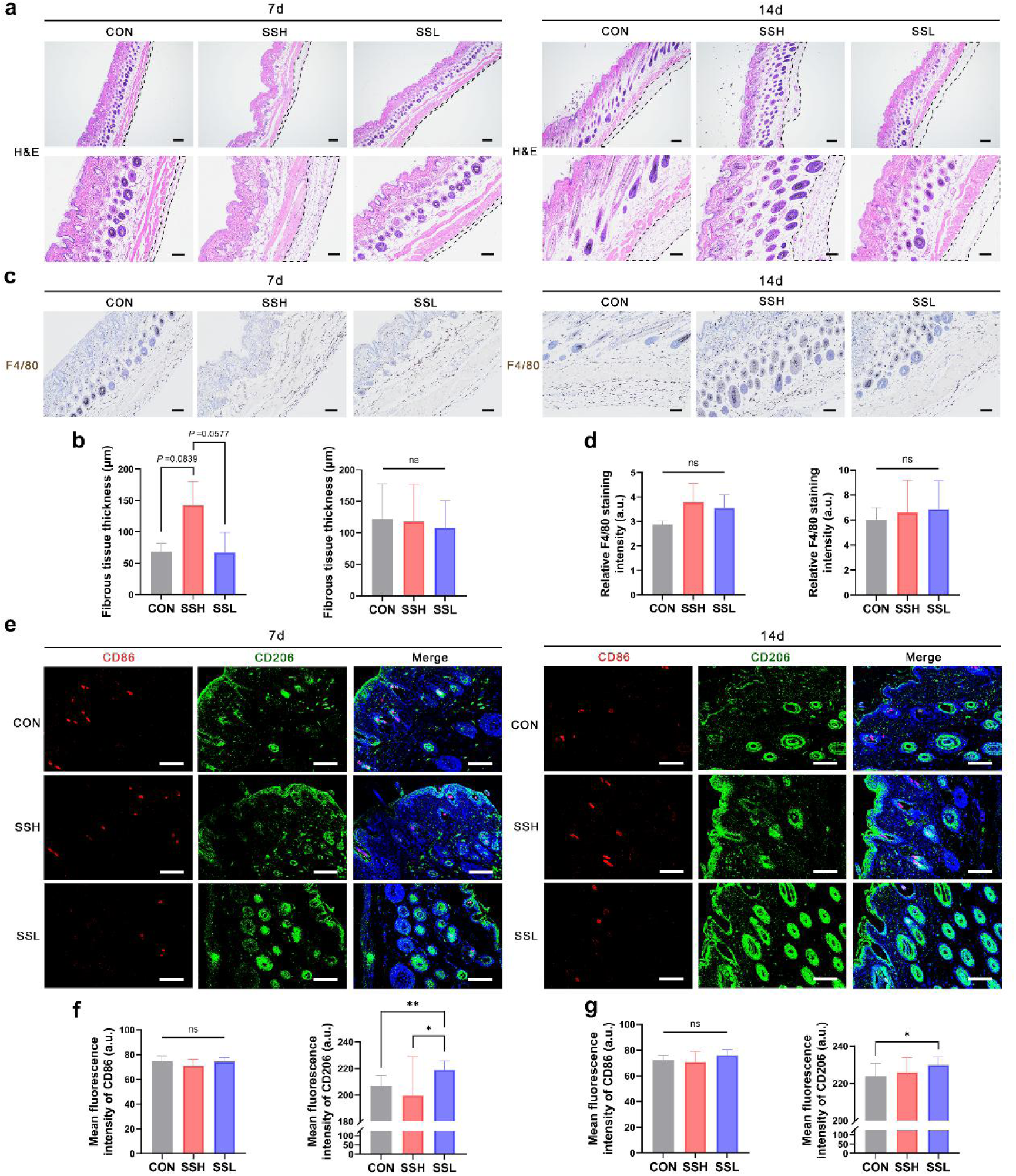
Inflammatory response of SSH and SSL in vivo. (a, b) H&E images on 7 and 14 days. The black dotted lines represent fibrous deposition. Scale bar, 200 μm in the upper panel and 100 μm in the lower panel. (c, d) Immunochemistry images and quantitative analysis by staining F4/80 macrophages on 7 and 14 days. (e-g) Immunofluorescent staining and quantitative analysis of CD86 and CD206 on 7 and 14 days. Scale bar, 100 μm. Data represent mean ± SD, n=6 mice per group, \**P* < 0.05, \*\**P* < 0.01, ns: not significant.

## 4. Discussion

Structurally, SSH extracted by boiling water displayed a nanofibrous morphology, whereas SSL appeared as nanoparticles. This distinction is likely due to their divergent secondary structures: SSH was β-sheet–dominant, while SSL exhibited a predominantly random-coil conformation, consistent with previous reports ^[12a, 15b]^. In the case of silk fibroin, both SFH and SFL formed nanoparticles and similarly adopted random-coil–rich structures. This observation aligns with earlier findings showing that degumming cocoons for 30 or 60 minutes yields fibroin fractions with different MW distributions but comparable secondary structures ^[13c]^. Furthermore, regarding the relationship between hydrodynamic radius and the molecular weights of SF, we did not detect the expected positive correlation, which may result from the spontaneous self-assembly behavior of fibroin molecules in aqueous environments ^[14b]^.

We found that SS with high- and low-MWs exerted opposite effects on the migration and tube-forming abilities of HUVECs. The fibrous network structure of SSH may partly account for its inhibitory effects on endothelial migration and tube formation. Furthermore, transcriptomic analysis indicated that the angiostatic properties of SSH may be attributed to the downregulation of *Id1* and *Smad6/9*, genes known to facilitate angiogenesis; inhibition of these genes has been reported to result in dysfunctional vasculature ^[17]^. In contrast, SSL upregulated the mRNA expression of angiogenesis-promoting genes such as *Samd9* and *Samd9l*, among others ^[18]^. These findings are consistent with a previous report showing that both SS and SF exhibit pro-angiogenic activity in HUVECs ^[8c]^. In our study, transcriptomic data further suggested that activation of the TGF-β signaling pathway may underlie the pro-angiogenic effects of SSL.

Our study demonstrated that SS with high- and low-MWs enabled macrophage-mediated angiogenesis in distinct ways. SSH induced a mixed M1/M2-like macrophage phenotype, whereas SSL primarily promoted an M2-like phenotype. Macrophages with M2-like characteristics are widely associated with pro-angiogenic activity ^[19]^, while pro-inflammatory macrophages can also secrete VEGF to stimulate angiogenesis ^[20]^. Summarizing previous studies on the interaction between SS and macrophages, SS has been reported to induce M1-like macrophages to produce VEGF ^[8d]^ or promote osteogenesis ^[15b]^, and, in other cases, to induce both M1- and M2-like macrophages to enhance angiogenesis ^[8a, 8b, 21]^—though most studies did not specify the MW of SS. In addition, we observed that SSH and SSL—differing in β-sheet content and morphology—elicited different macrophage polarization patterns. This raises the possibility that the structure of SS may contribute to functional outcomes. A previous study showed that SS with molecular weights above 30 kDa and higher β-sheet content drove M1-like macrophages to induce osteogenesis ^[15b]^. However, in our study, we do not have sufficient evidence to establish a direct link between β-sheet content (or MW) and macrophage polarization, as we detected a trace amount of endotoxin (<0.2 EU/mL) in SSH. Although this level is below the regulatory limit typically considered acceptable (0.5 EU/mL) ^[22]^, we cannot exclude the possibility that endotoxin contamination contributed to the M1-like phenotype observed in SSH. In addition, the angiogenic effects observed in our conditioned media experiments likely arise from a complex mixture of soluble cytokines, growth factors, and possibly extracellular vesicles.

For SFH and SFL, no marked differences were observed in their direct effects on HUVEC angiogenesis. However, their indirect effects through macrophages differed: SFH did not noticeably influence macrophage polarization, whereas SFL induced an M2-like phenotype that may promote angiogenesis. This suggests that SFL possesses higher bioactivity than SFH. Previous studies have similarly reported that low-MW SF or SF-derived peptides can enhance cell proliferation ^[15d]^ and modulate various cellular responses^[15e]^. Considering that SF-based biomaterials undergo gradual degradation in vivo, resulting in decreased MWs over time, such MW-dependent changes may help explain the angiogenic activity frequently observed in SF-based biomaterials.

## 5. Conclusions

In summary, our study highlights the critical role of molecular weight in modulating the angiogenic properties of silk proteins, particularly silk sericin. Variations in molecular weight not only alter the structural characteristics of silk proteins but also lead to marked differences in their biological functions. Reducing molecular weight shifts the secondary structures of the silk proteins from β-sheet–rich to random coil–dominant conformations. High-molecular-weight silk sericin (40-180 kDa) induces a mixed M1/M2-like macrophage response, promoting angiogenesis but simultaneously compromising vascular integrity and increasing fibrosis. In contrast, low-molecular-weight silk sericin (less than 25 kDa) drives M2-like macrophage polarization, and facilitates angiogenesis along with enhanced vascular maturation and reduced fibrotic reactions. These findings not only deepen our understanding of silk protein–mediated bioactivities but also provide insight into how molecular weight tuning may be leveraged in other biopolymers to guide biomaterial design for tissue engineering and regenerative medicine.

## Acknowledgments

This work was supported by the Guangdong Basic and Applied Basic Research Foundation (2022A1515110329).

## Data availability

Data will be made available on request.

## Conflict of interest

The authors declare no conflict of interest.

## Abbreviations

SS: silk sericin
SF: silk fibroin
SSH: high molecular weight silk sericin
SSL: low molecular weight silk sericin
SFH: high molecular weight silk fibroin
SFL: low molecular weight silk fibroin
CON: control group
HUVEC: human umbilical vein endothelial cell
TNF-α: tumor necrosis factor-α
VEGF: vascular endothelial growth factor

## Notes

### Competing Interest Statement

The authors have declared no competing interest.

