## supporting info for "Angiogenesis Guided by *Bombyx mori* Silk Proteins is Molecular Weight-Dependent"

Tiandong Li, Jian He, Yujian Jiang, Jiajun Chen, Yao Xu, Yeyuan Wang, Jingchen Sun*, Doudou Hu*

Guangdong Engineering Technology Research Center of Sericulture, College of Animal Science, South China Agricultural University, Guangzhou, Guangdong, 510642, China

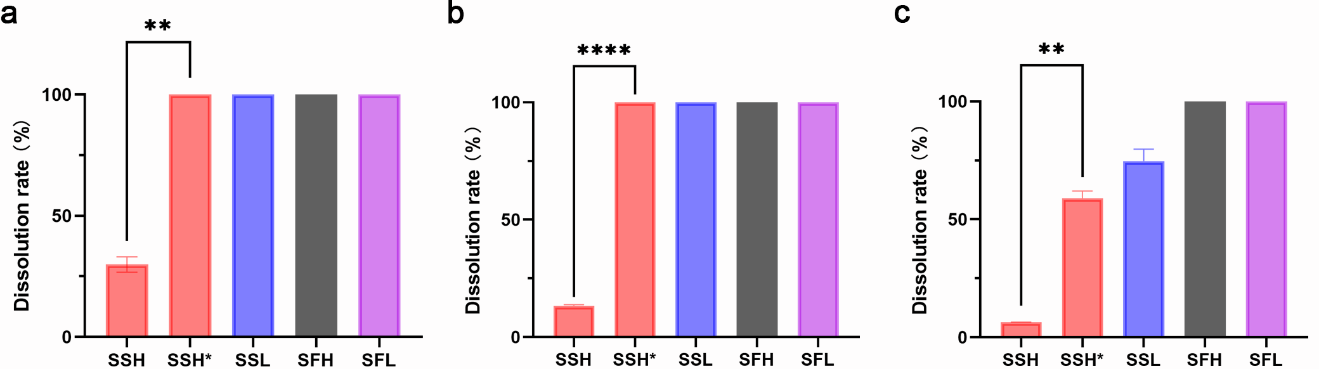
Figure S1. Solubility of different silk proteins. The concentration was (a) 0.1% (w/v), (b) 0.5% (w/v), and (c) 1% (w/v). SSH*: SSH heated by boiling water bath for 10-15 min. Data represent mean ± SD, n=3, ***P* < 0.01, *****P* < 0.0001.

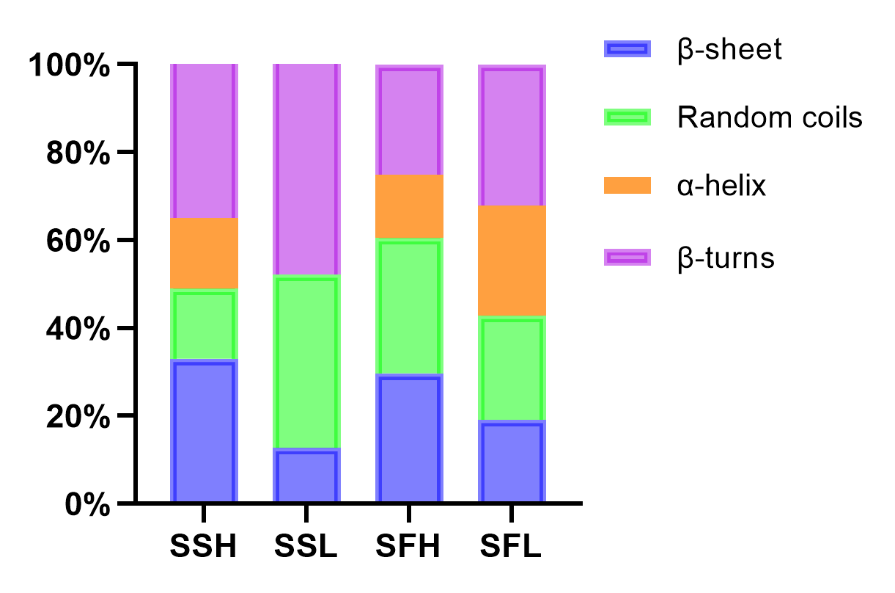

Figure S2. The secondary structure percentage of the four silk proteins calculated by deconvolution of the FTIR spectra.

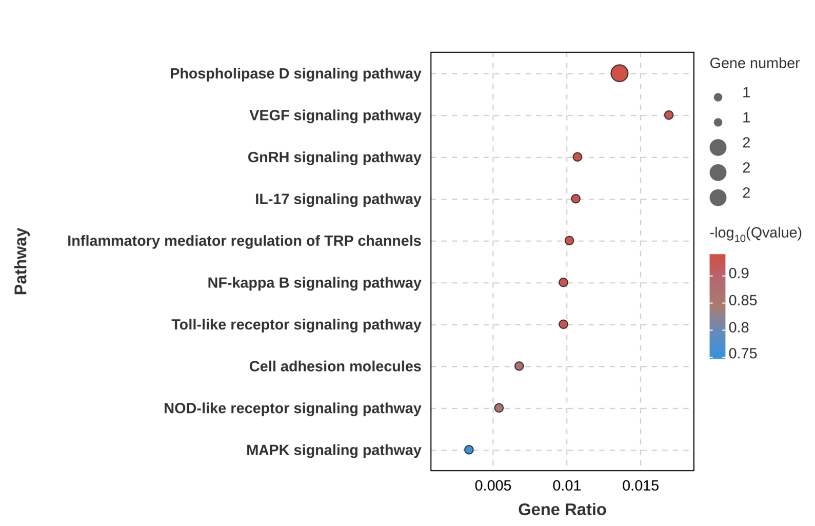

Figure S3. KEGG analysis of different signaling pathways between CON and SFH.

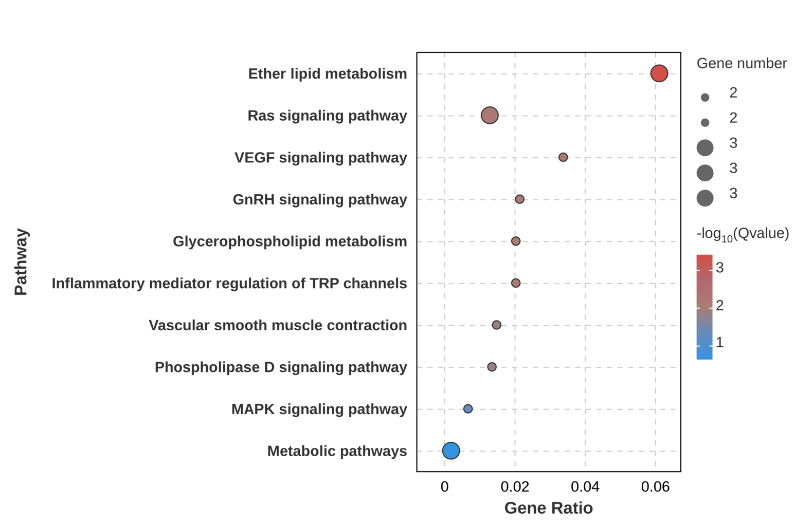

Figure S4. KEGG analysis of different signaling pathways between CON and SFL.

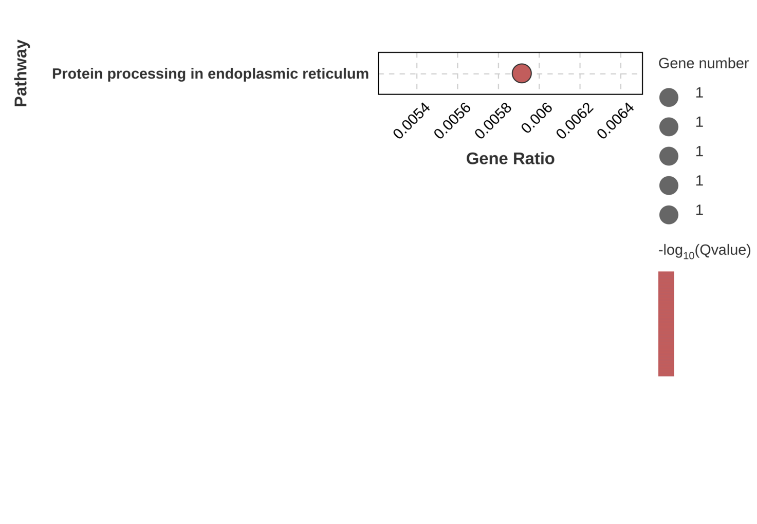

Figure S5. KEGG analysis of different signaling pathways between SFH and SFL.

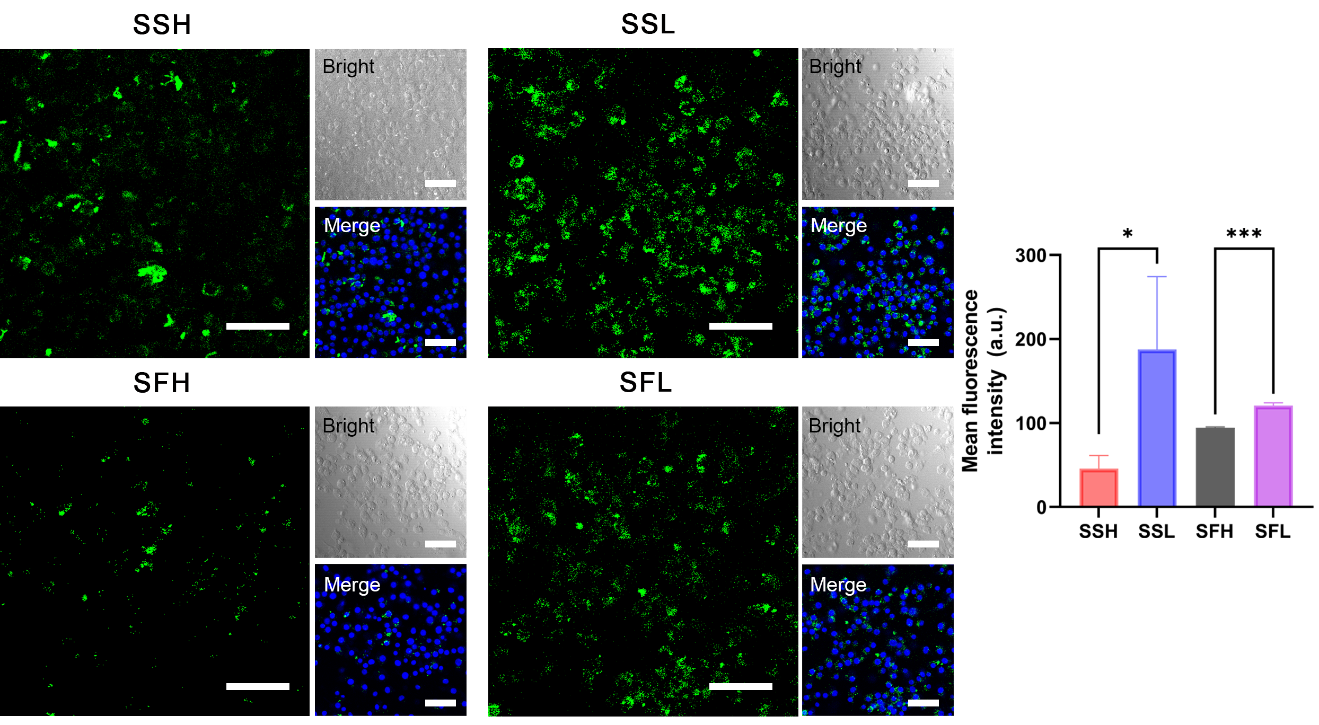

Figure S6. Confocal images and quantitative analysis of cellular uptake in RAW 264.7 cells treated with 0.01% (w/v) FITC-labeled silk proteins (green) for 24 h. Scale bar, 50 μm. Data represent as mean ± SD, n=3. **P* < 0.05, ****P* < 0.001.

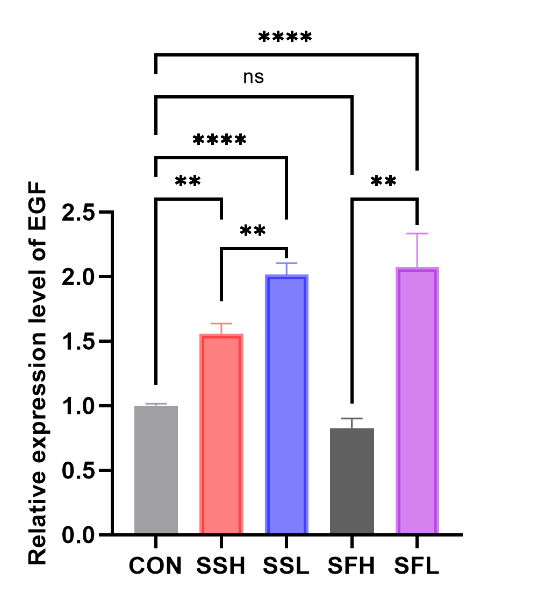

Figure S7. RT-qPCR analysis of EGF secreted by RAW264.7 cells treated with 0.01% (w/v) silk proteins for 24 h. Data represent as mean ± SD, n=3. **P* < 0.05***P* < 0.01, ****P* < 0.001, *****P* < 0.0001, ns: not significant.

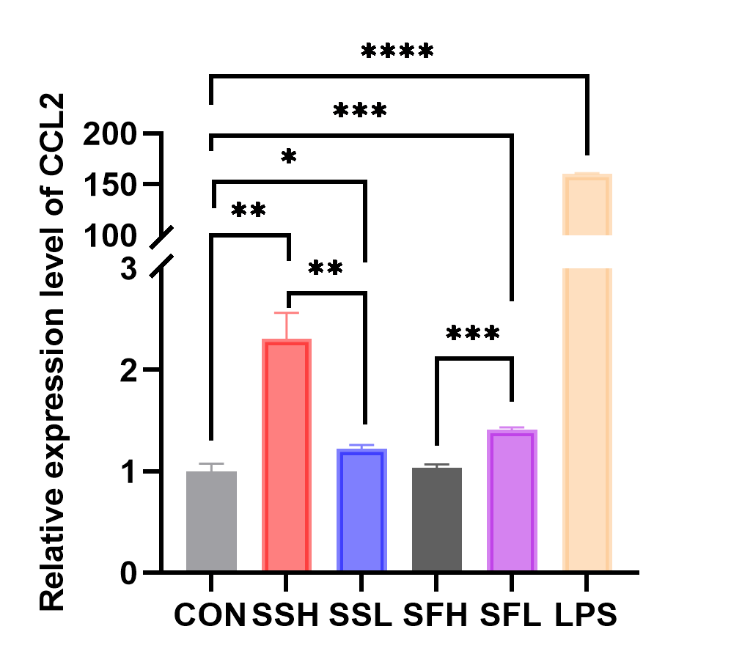

Figure S8. RT-qPCR analysis of CCL2 in HUVECs treated with conditioned medium from silk-treated RAW 264.7 cells. Data represent as mean ± SD, n=3. **P* < 0.05, ***P* < 0.01, ****P* < 0.001, ****P* < 0.001.

Table S1. List of primers for qPCR analysis.

| **Primers** | **Forward primer sequence F: (5'-3')** | **Reverse primer sequence R: (5'-3')** | **species** |
| --- | --- | --- | --- |
| GAPDH | TGACGCTGGGGCTGGCATTG | GGCTGGTGGTCCAGGGGTCT | Human |
| VEGFA | CTTGCCTTGCTGCTCTACC | CACACAGGATGGCTTGAAG | Human |
| CCL2 | CAGCCAGATGCAATCAATGCC | TGGAATCCTGAACCCACTTCT | Human |
| EGF | CACTTCCGCTTGGCTCATCA | CATCATGGTGGTGGCTGTCTG | Mouse |
| GAPDH | TTCCAGGAGCGAGACCCCACTA | GGGCGGAGATGATGACCCTTTT | Mouse |
